# Genetic characterization of extended-β-lactamase (ESBL) plasmids captured from dairy manures

**DOI:** 10.1101/2023.03.20.533445

**Authors:** Tam Tran, Joe Thorne, Andrew Scott, James Robertson, John HE Nash, Catherine Carrillo, Edward Topp

## Abstract

This study was to assess the gene diversity and characterize a large set of plasmids harboring extended β-lactamase (ESBL) genes from raw and digested dairy manure. A total of eighty-four plasmids that were captured in this *E. coli* recipient were sequenced using Illumina MiSeq sequencing technology. Twenty-four plasmids of interest were subsequently sequenced using MinION technology in order that a hybrid assembly could be performed on short-and long-read sequences to circularize and complete these plasmids. The size of sequenced plasmids ranged between 40 and 260 kb with various incompatibility groups: IncC, IncI1, IncN, IncY, IncB/O/K/Z, IncX1, IncHI2, IncHI2A, IncFIB(K), IncFII. A variety of extended β-lactamase genes were identified: *bla*_CTXM -1_, *bla*_CTXM -14_, *bla*_CTXM -15_*, bla*_CTXM-27_*, bla*_CTXM-55_, *bla*_CTXM-61_, *bla*_PER-1,_ *bla*_IMP-27_. Interestingly, the *bla*_IMP-27_ gene, a novel metallo-β-lactamase discovered in the last decade, was found located on an integrated region in the host chromosome. And one plasmid carrying the *bla*_CMY-2_ gene, an AmpC gene, also expressed ESBL phenotype. Four virulence factors, including *cia, cib, traT* and *terC,* were detected on some of these plasmids. In addition, six type-2 toxin-antitoxin systems were detected: MazF/E, PemK/I, HipA/B, YdcE/D, RelB/E and HigB/A. Twenty-two out of twenty-four complete plasmids carried putative prophage regions; and most of prophage hits were marked as incomplete, except that the largest plasmid pT525A and the IncY plasmid pT415A had prophage hits with higher scores.

**IMPORTANCE:** The widespread of antibiotic resistant bacteria is largely due to the exchange of mobile genetic elements such as plasmids. Plasmids harboring extended β-lactamase (ESBL) genes originated from dairy manure potentially become entrained in manured soil, which subsequently enter the human food chain. Currently there is a lack of detailed information on these plasmids in the environment, specifically in dairy manure. This study unveils the abundance and diversity of ESBL-carrying plasmids from both raw and digested manures which were captured in *gfp-*labelled *E. coli* CV601. In addition, the study provides insightful information of plasmid characteristics including incompatibility groups, ESBL genes combined with other resistance genes, mobile genetic elements (transposons, insertion sequence), toxin-antitoxin systems, virulence factors and prophage sequences.

## INTRODUCTION

Extended-β-lactamase (ESBL) genes have been a matter of undoubtedly grave public-health concern due to their ability to hydrolyze third-generation cephalosporins (e.g. cefotaxime, ceftriaxone, ceftazidime, or cefepime) and monobactams (aztreonam) (1). A dramatic increase in the number of multidrug-resistant Enterobacteriaceae (mostly *Escherichia coli*) that produce extended-spectrum β-lactamases (ESBLs), such as the CTX-M enzymes, has been reported since the 1990s (2). ESBL genes has been widely disseminated via mobile genetic elements such as plasmids, insertion sequences, transposons. Plasmids carrying ESBL genes are ubiquitous in environments including manure, manured soil, wastewater treatment plants and aquaculture (3–8).

Bacterial toxin-antitoxin (TA) systems are pairs of genes encoding a toxin protein and its corresponding antitoxin protein which can be found on either chromosomes or plasmids in free-living bacteria (9, 10). The first TA operon was found on plasmid R1 about three decades ago, and was shown to play an important role in plasmid stability by the post-segregational removal of plasmid-free cells (11, 12). The ccd system on the F plasmid, the most widely studied system, was even employed in DNA cloning strategies (13). Depending on the molecular structures and mechanisms of action, three types of TA operon were presented: Type I, II, and III (12). The type II TA system, also termed as the addiction system, consisted of at least ten current families such as MazE-MazF, RelE-RelB, YefM-YoeB, and MqsR-MqsA (12, 14). Despite their ubiquity in bacteria, TA systems on manure-originated plasmids are not well-understood.

Virulence factors mainly accounts for bacterial pathogenicity which causes diseases in hosts such as plant, animals and human (15, 16). They can be found on either pathogenicity islands in the genome of pathogenic bacteria or on plasmids (17, 18). Virulence-associated plasmids in *E. coli* were associated with six pathotypes enterotoxigenic *E. coli* (ETEC), enteroaggregative *E. coli* (EAEC), enteroinvasive *E. coli* (EIEC), enterohemorrhagic *E. coli* (EHEC), enteropathogenic *E. coli* (EPEC), extraintestinal pathogenic *E. coli* (ExPEC) (17). Nine out of 26 plasmid incompatibility groups which mostly fall into group F were known to carry virulence genes, and there is no doubt this number would keep rising as novel plasmid groups continue to be identified (17, 19).

Prophages are bacteriophage sequences that normally integrate into bacterial chromosome and largely contribute to bacterial adaptation and evolution by enabling the horizontal genetic exchange (20, 21). A few prophages (e.g. P1, N15, LE1, ɸ20, and ɸBB-1), however, are able to independently replicate in the lysogen as low-copy-number plasmids (22). Prophages have an average size between 29 to 78 kb, which probably constitute about 0.6 to 1.8 % of the host chromosome (23). Therefore, plasmids of megasize (> 100kb) can easily capture prophage regions via either homologous recombination or the movement of insertion sequences/ transposons. Recent evidences suggested that the plasmid pMCR-1-P3, an IncY plasmid, was the outcome of homologous recombination event between a plasmid and a prophage region located in the *E. coli* genome (24).

The aim of the study was to extensively and intensively analyze genetic characteristics of eighty-three sequenced ESBL plasmids originated from dairy manure including a subset of twenty-five plasmids reported previously (25). The study also revised the comparison of plasmids from raw with those from digested manure in a larger set of data. Overall this study provide insightful information of plasmid characteristics such as plasmid size, virulence factor, TA systems, incompatibility groups, mobile genetic elements and antibiotic resistance genes

## RESULTS

### Description of sequenced plasmids harboring ESBL genes from raw and digested manures

In this study, a total of 83 plasmids harboring ESBL genes were grouped based on their incompatibility groups, ESBL genes and other resistance genes (Table 1). A detailed information of 25 plasmids was reported previously (manure paper); while the rest of them can be found in supplementary material of this study (Table S1). Twenty distinct plasmid profiles carrying eight ESBL genes in combination with other resistance genes were identified (Table 1). Eleven transconjugants’ whole genomes were further sequenced on the MinION long-read sequencing platform so that hybrid assembly could be used to completely close the plasmids carried by these transconjugants. Maps of these complete plasmids were presented in Fig. 1 & Fig. S1.

**Fig. 1:**
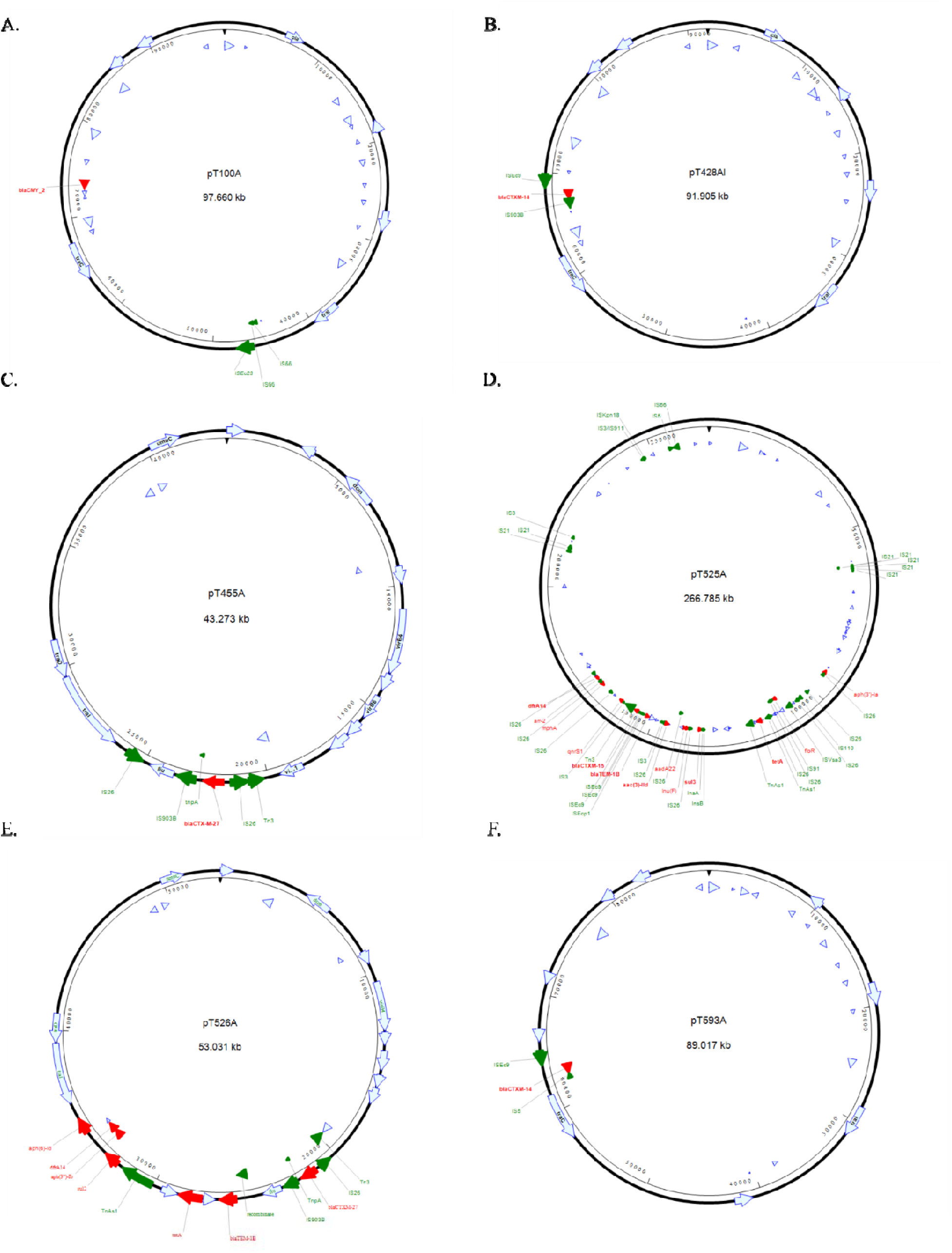
Plasmid maps of six distinct plasmids harboring ESBL/AmpC genes which were captured in *E. coli* CV601 strain. (A) pT100A, (B) pT428Al, (C) pT455A, (D) pT525A, (E) pT526A, (F) pT593A. Red arrows are resistance genes detected by starAMR tool. Green arrows are mobile genetic elements detected by RAST and BLAST tools. Dark blue arrows are other functional genes which were annotated by PROKKA tool.

**Table 1:**
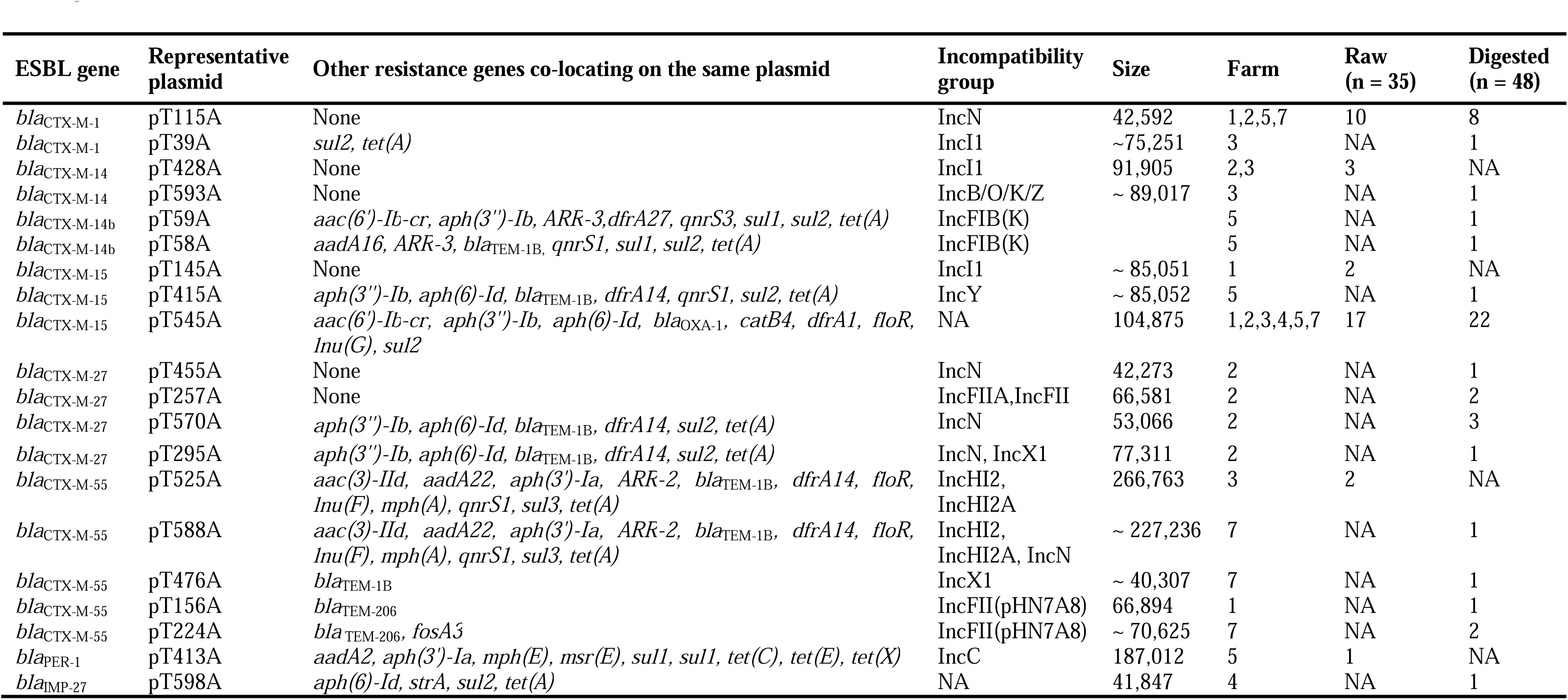
Distribution of eighty-three sequenced plasmids based on their incompatibility groups, ESBL genes and other resistance genes

Among 83 sequenced plasmids, 35 of them were from raw manure, and 48 from digested manure (Table 1). The most frequent plasmid (pT545A) carried *bla*_CTXM-15_ gene along with other nine resistance genes: *aac(6’)-Ib-cr, aph(3’’)-Ib, aph*(*6*)*-Id, bla*_OXA-1_*, catB4, dfrA1, floR, lnu(G), sul2*. This plasmid was found in both raw and digested manures from all participating farms. The second most frequent plasmid carrying *bla*_CTXM-1_ gene (pT115A) was found in both raw and digested manure from four out of six participating farms.

Other 18 less frequent plasmids were found in either raw (four plasmids) or digested manures (fourteen plasmids) (Table 1). Four plasmids from raw manure individually carried following ESBL genes: *bla*_CTX-M-14_, *bla*_CTX-M-15_, *bla*_CTX-M-55_, *bla*_PER-1_. Meanwhile, thirteen out of fourteen plasmids from digested manure individually carried following ESBL genes: *bla*_CTX-M-1_, *bla*_CTX-M-14_, *bla*_CTX-M-14b_, *bla*_CTX-M-15_, *bla*_CTX-M-27_, *bla*_CTX-M-55_. Interestingly, one conjugative plasmid (pT598A) from digested manure did not carry any ESBL genes. However, the *bla*_IMP-27_ gene was detected on the host chromosome along with other resistance genes: *aph*(*6*)*-Id, strA, sul2, tet(A)* in the transconjugant carrying this plasmid pT598A.

Sizes of hybrid assembled plasmids ranged between 40 and 260 kbs; the most frequent plasmid (pT545A) had the size of 100 kbs. Incompatibility groups identified in this study were IncC, IncI1, IncN, IncY, IncB/O/K/Z, IncX1, IncHI2, IncHI2A, IncFIB(K), IncFII(pHN7A8). A variety of mobile genetic elements that were located in areas surrounding resistance genes were identified including Tn*3*, Tn*7*, Tn*As1*, *tnpA*, *IntI1*, *insA*, *insB*, IS*Ec63*, IS*Ec9*, IS*Ecp1,* IS*903B*, IS*3*, IS*5*, IS*26*, IS*91*, IS*110*, IS*5075*, and IS*Vsa3*. Mobile genetic elements that were not located proximal to resistance genes were also identified: IS*3*, IS*5*, IS*66*, IS*21*, IS*911*, IS*Kpn18*, IS*Ec23*.

### Description of plasmids with special features

The plasmid pT100A was the only conjugative plasmid carrying an AmpC-type gene (*bla*_CMY-2_); however, the transconjugant carrying this plasmid expressed ESBL phenotype (Fig 1. & Table S1). The plasmid with a size of 98 kbs had IncI1 incompatibility group. The plasmid did not carry any other resistance genes, but it carried two conjugal transfer genes (*traC*, *traI*). Unlike most of captured plasmids, mobile genetic elements on this plasmid were located far apart from the resistance gene *bla*_CMY-2_.

The plasmid pT413A with a size of 187 kbs carrying the *bla*_PER-1_ gene, an ESBL gene, also carried mercury resistance operon (Fig. 2). The mercury resistance operon with a size of approximately 3 kbs consisted of *merA*, *merP*, *merT* and *merR* genes. This operon located in the 79 kb region along with other resistance genes and other mobile genetic elements (integrons, transposon and insertion sequences). The adjacent mobile genetic elements surrounded this operon were transposon Tn7 transposition proteins (TnsB, TnsC).

**Fig. 2:**
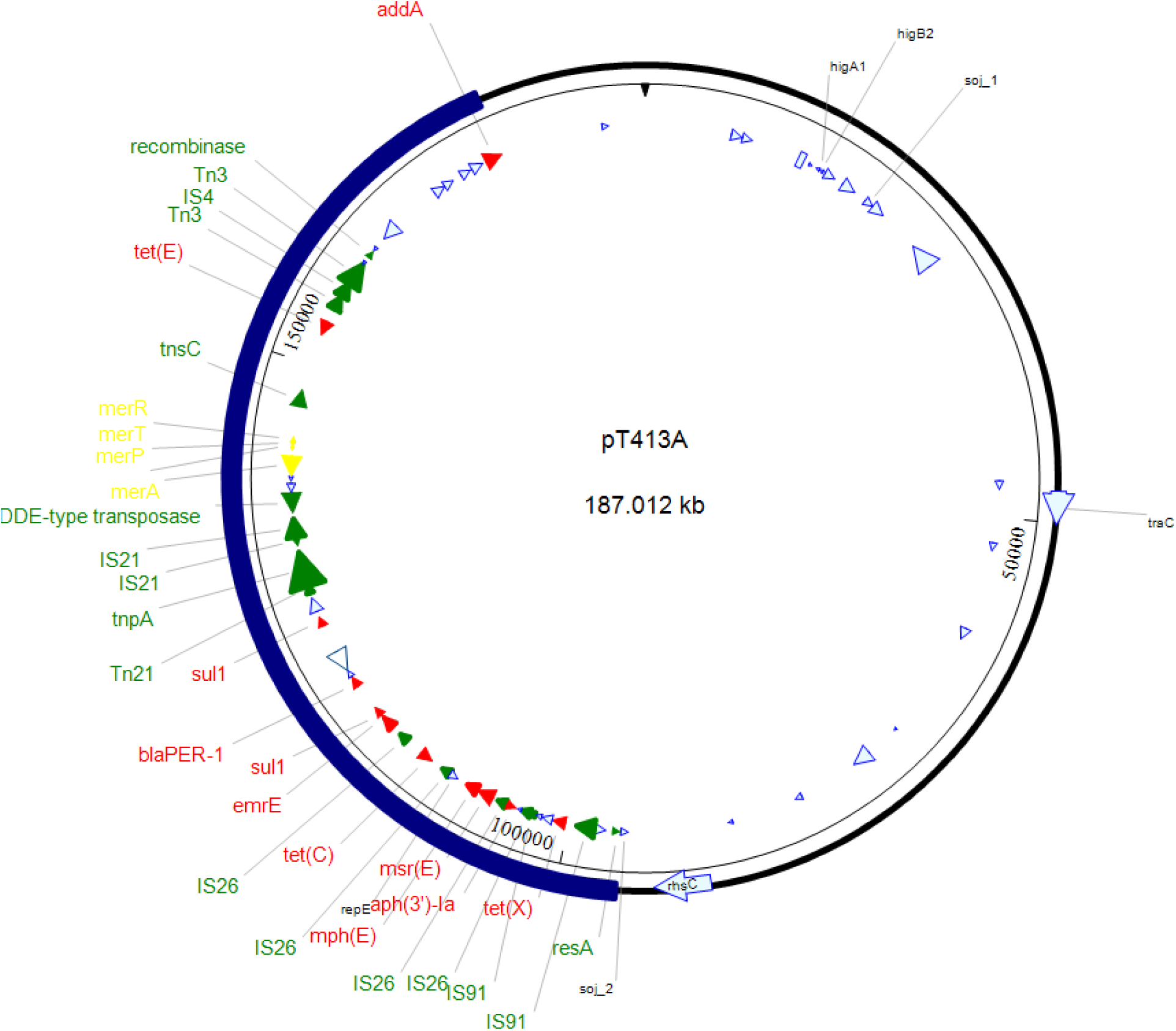
The map of plasmid pT413A (187 kb). The blue stripe indicates the 79 kb-region harboring mobile genetic elements (green arrows and labels), mercury resistance operon (yellow arrows and labels) and antibiotic resistance genes (red arrows and labels).

The conjugative plasmid pT428Al carrying the *bla*_CTX-M-14_ gene accompanied with the mobilizable plasmid pT428As which did not carry any resistance genes. The pT428Al had a size of 92 kbs, and its accompanying plasmid pT428As had a size of 4kbs. This is the only case where two plasmids were confirmed to co-transfer into *E. coli* recipient CV601 by hybrid assembly. Using the annotation tool PROKKA, only two genes, *mobA* encoding mobilization protein A and *repE* encoding replication initiation protein, were detected on the smaller plasmid pT428As.

The plasmid pT598A was the only conjugative plasmid did not carry any resistance genes. However, this plasmid carried many conjugal transfer genes: *traB, traC, traD, traG, traI, traK, traJ, traL, traM.* The plasmid pT598A with a size of 42 kbs had no incompatibility group identified. Hybrid assembly of transconjugant’s whole genome sequence carrying this plasmid showed a 865kb region integrated into the host chromosome (Fig. 3). This region carried multiple resistance genes (*aph*(*6*)*-Id, bla*_IMP-27_*, strA, sul2, tet(A)*) along with other mobile genetic elements (Tn*As1*, IS*Vsa3*, IS*3*, IS*26*, *intA*). The integrated region was between two chromosomal genes: *aspC* gene encoding aspartate aminotransferase and *asnS* gene encoding asparagine--tRNA ligase.

**Fig. 3:**
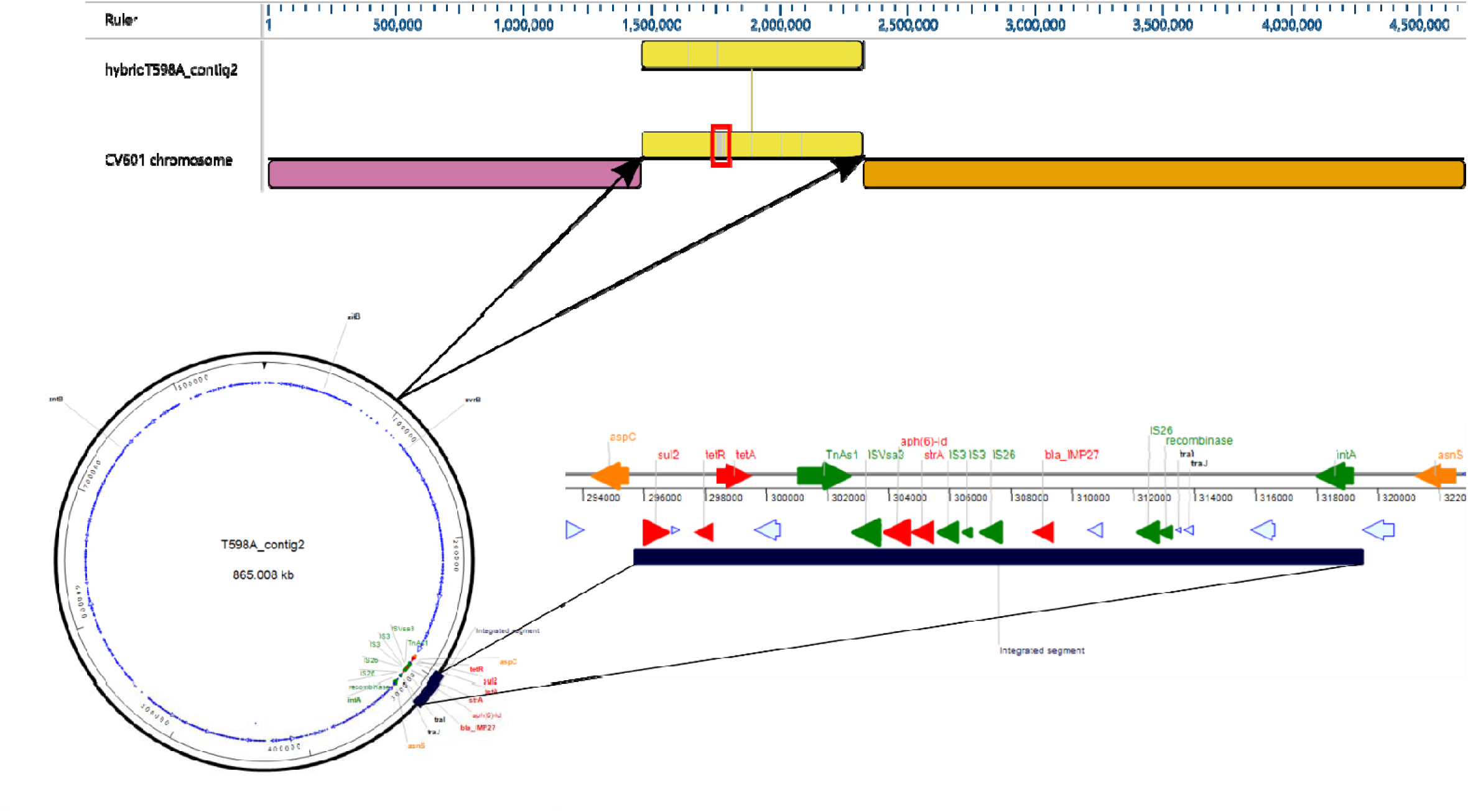
Map of an integrated region containing multiple mobile genetic elements and resistance genes including the metallo-β-lactamase gene *bla*_IMP-27_. The top figure shows the alignment of the contig #2 of the isolate T598A with the host chromosome *E. coli* CV601; the red box indicates the location where this *bla*_IMP-27_-bearing region got integrated. The bottom left figure shows the circular form of contig #2 with the integrated region. The bottom right figure is the enlarged integrated region. Red labels/arrows are resistance genes; green labels/arrows are mobile genetic elements (transposon, insertion sequence, integron); orange labels/arrows are genes on the host chromosome adjacent to the integrated region; blue arrows are other functional genes.

### DNA variations among genotypically similar plasmids

In this study, not all plasmids were sequenced on both short-read and long-read sequencing platforms. Plasmids that were sequenced on both platforms were successfully completely closed via hybrid assembly, hence they could be used as a reference input. Plasmids having similar characteristics (sizes, resistance genes, incompatibility groups) with closed plasmids were further analyzed using Snippy tool to search for SNPs or any DNA variations.

There were very few DNA variation detected among plasmids harboring the *bla*_CTX-M-1_/ *bla*_CTX-M-15_/ *bla*_CTX-M-27_ gene (Table S2, S3 and S4). However, three largest IncHI2-IncHI2A plasmids (>200 kb) harboring the *bla*_CTX-M-55_ gene were quite distinct from one another (Table S5). Unexpectedly, plasmid pT525A was even more different from plasmid pT594A considering they both were originated from raw manure of the same farm, and they had resistance genes and incompatibility groups in common.

**Table 2.**
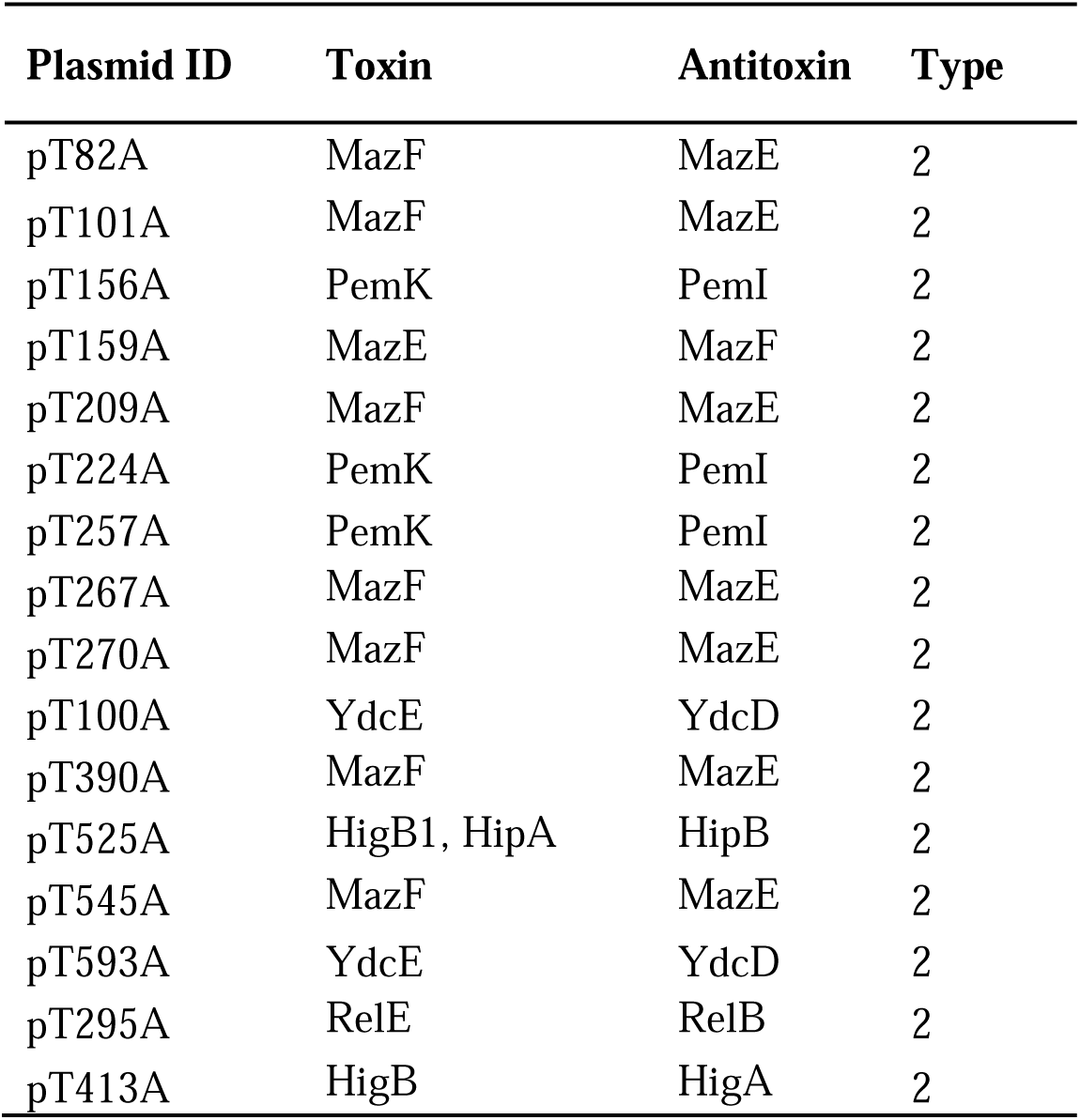
List of toxin-antitoxin systems found on plasmids

| Plasmid ID | Toxin | Antitoxin | Type |
| --- | --- | --- | --- |
| pT82A | MazF | MazE | 2 |
| pT101A | MazF | MazE | 2 |
| pT156A | PemK | PemI | 2 |
| pT159A | MazE | MazF | 2 |
| pT209A | MazF | MazE | 2 |
| pT224A | PemK | PemI | 2 |
| pT257A | PemK | PemI | 2 |
| pT267A | MazF | MazE | 2 |
| pT270A | MazF | MazE | 2 |
| pT100A | YdcE | YdcD | 2 |
| pT390A | MazF | MazE | 2 |
| pT525A | HigB1, HipA | HipB | 2 |
| pT545A | MazF | MazE | 2 |
| pT593A | YdcE | YdcD | 2 |
| pT295A | RelE | RelB | 2 |
| pT413A | HigB | HigA | 2 |

### Toxin-antitoxin systems detected on completely closed plasmids

Six type-2 toxin-antitoxin systems were detected: MazF/E, PemK/I, HipA/B, YdcE/D, RelE/B and HigB/A (Table 2). The MazF-MazE system were the most frequent toxin-antitoxin system detected in our study. It was detected on following plasmids: pT82A, pT101A, pT159A, pT209A, pT267A, pT270A, pT390A, pT545A. The PemK-PemI system was found located on three plasmids: pT156A, pT224A and pT257A. These three plasmids shared a majority of their sequence in common (25). The YcdE-YcdD system were detected on two distinctly different plasmids pT100A and pT593A. The remaining three systems found on less frequent plasmids: HipA-HipB system on pT525A, RelE-RelB system on pT295A and HipA-HipB on pT525A. There was no toxin-antitoxin systems detected on following plasmids: pT199A, pT247A, pT115A, pT428Al, pT428As, pT455A, pT526A, pT570A.

### Detection of virulence factor genes on completely closed plasmids

There were four virulence factors detected: *cia, cib, traT, terC* (Table 3). The *cia* gene encoding colicin ia was found on two distinct plasmids pT247A and pT428Al, while the *cib* gene encoding colicin ib was detected on one plasmid pT100A. The *traT* gene encoding complement resistance protein was found on four plasmids pT156A, pT224A, pT257A and pT593A. Among them, pT593A was more distinctly different than the other three plasmids (pT156A, pT224A, pT257A) whose sequences shared a lot in common as shown previously (25). The *terC* gene encoding tellurium ion resistance protein was detected on one plasmid pT525A. There was no virulence factor detected on following plasmids: pT82A, pT100A, pT101A, pT115A, pT159A, pT209A, pT267A, pT270A, pT295A, pT390A, pT413A, pT428As, pT545A, pT526A, pT570A, pT593A, pT598A, There was no shiga-toxin genes was detected on any of plasmid input.

**Table 3.** Detection of virulence factors on plasmids

| Plasmid ID | Virulence factor | Identity | Query / Template length | Position in contig | Protein function | Accession number |
| --- | --- | --- | --- | --- | --- | --- |
| pT100A | *cib* | 100 | 1881 / 1881 | 6021..7901 | Colicin ib | KP198616 |
| pT428l | *cia* | 100 | 1881 / 1881 | 5247..7127 | Colicin ia | UDDL01000017 |

Table 3. Detection of virulence factors on plasmids
|  |  |  |  |  |  |  |
| --- | --- | --- | --- | --- | --- | --- |
| pT593A | *traT* | 100 | 627 / 627 | 51274..51900 | Outer membrane protein<br>complement resistance | MF156268 |
| pT525A | *terC* | 100 | 1041 / 1041 | 78902..79942 | Tellurium ion resistance protein | KU341381 |
| pT156A | *traT* | 100 | 732 / 732 | 53990..54721 | Outer membrane protein<br>complement resistance | CYCV01000028 |
| pT224A | *traT* | 100 | 734 / 735 | 56998..57732 | Outer membrane protein<br>complement resistance | NC_019073 |
| pT257A | *traT* | 100 | 735 / 735 | 52276..53010 | Outer membrane protein<br>complement resistance | NZ_CP032205 |
| pT247A | *cia* | 100 | 1881 / 1881 | 5247..7127 | Colicin ia | UDDL01000017 |

### Detection of prophage sequences on plasmids

A majority of plasmids (22/26) got hits for prophage detection (Table 4): three plasmids got three prophage hits (pT82A, pT390A, pT545A); seven got two prophage hits (pT100A, pT101A, pT247A, pT270A, pT428Al, pT525A, pT593A); 13 got one prophage hit (pT115A, pT156A, pT159A, pT199A, pT209A, pT224A, pT257A, pT267A, pT295A, pT415A, pT455A, pT526A, pT570A). Three plasmids did not have any prophage sequences detected: pT428As, pT598A, pT413A.

**Table 4.**
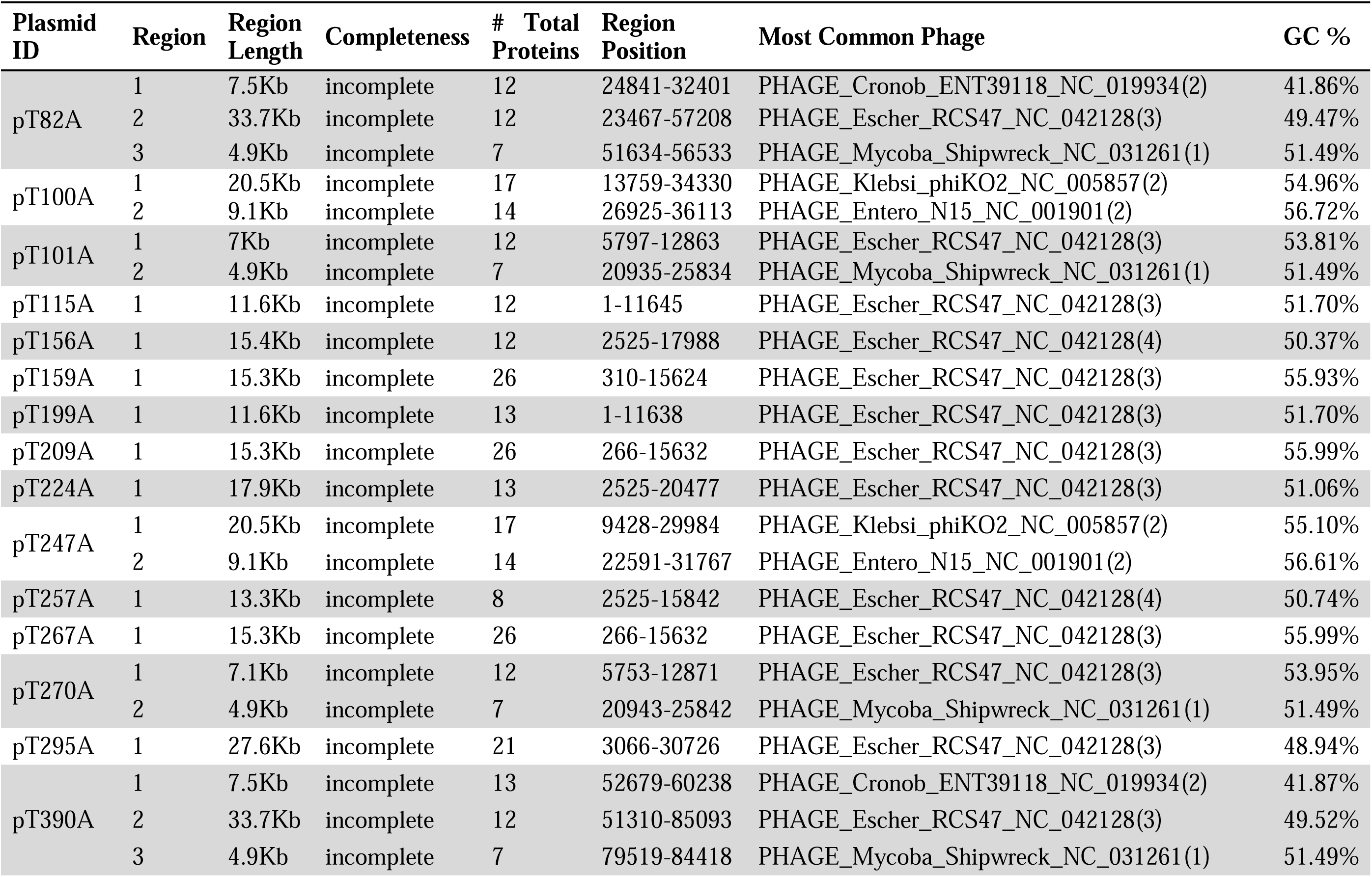
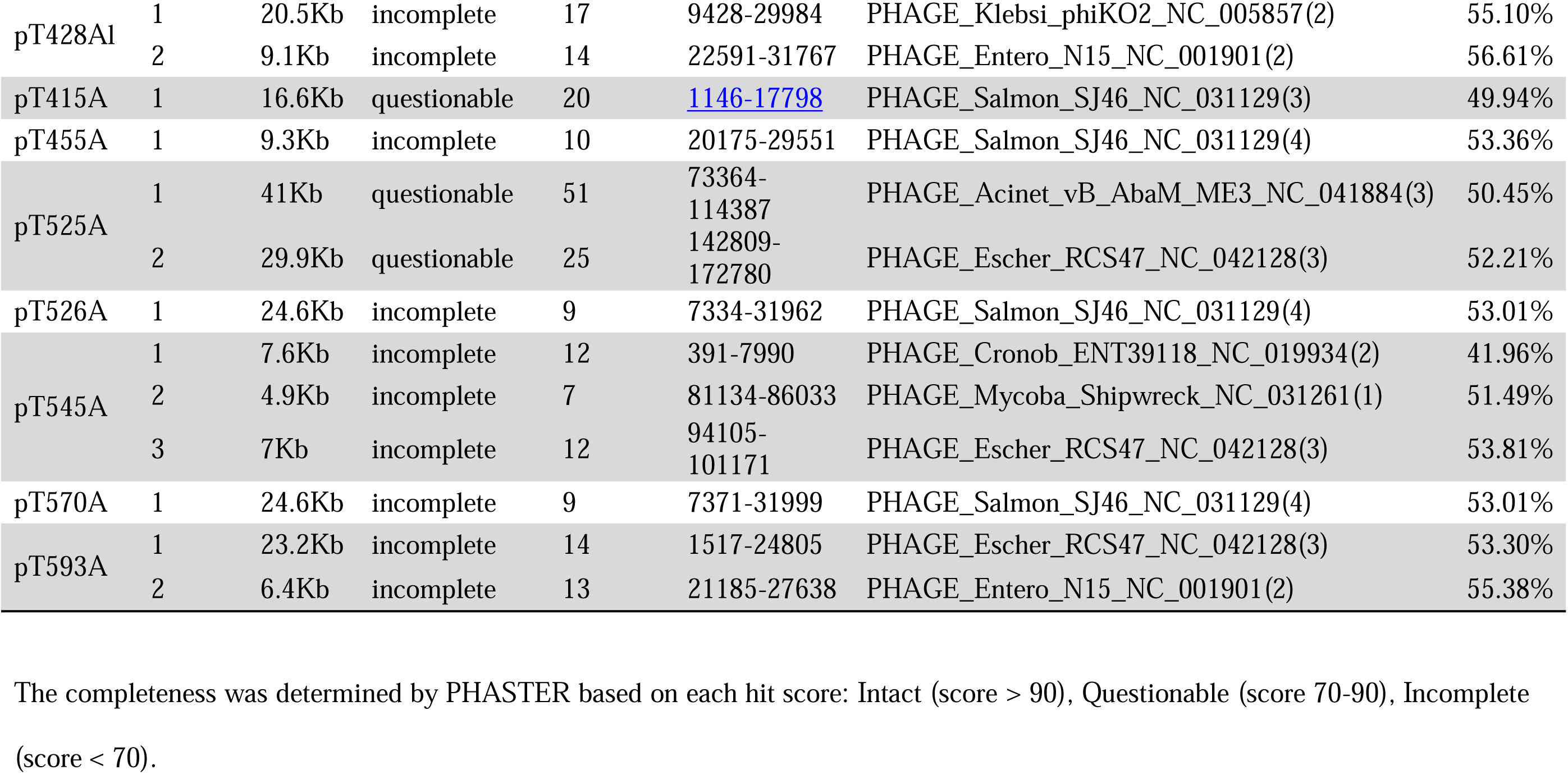
Identification of putative prophages on plasmids by PHASTER

| Plasmid ID | Region | Region Length | Completeness | # Total Proteins | Region Position | Most Common Phage | GC % |
| --- | --- | --- | --- | --- | --- | --- | --- |
| pT82A | 1 | 7.5Kb | incomplete | 12 | 24841-32401 | PHAGE_Cronob_ENT39118_NC_019934(2) | 41.86% |
|  | 2 | 33.7Kb | incomplete | 12 | 23467-57208 | PHAGE_Escher_RCS47_NC_042128(3) | 49.47% |
|  | 3 | 4.9Kb | incomplete | 7 | 51634-56533 | PHAGE_Mycoba_Shipwreck_NC_031261(1) | 51.49% |
| pT100A | 1 | 20.5Kb | incomplete | 17 | 13759-34330 | PHAGE_Klebsi_phiKO2_NC_005857(2) | 54.96% |
|  | 2 | 9.1Kb | incomplete | 14 | 26925-36113 | PHAGE_Enterо_N15_NC_001901(2) | 56.72% |
| pT101A | 1 | 7Kb | incomplete | 12 | 5797-12863 | PHAGE_Escher_RCS47_NC_042128(3) | 53.81% |
|  | 2 | 4.9Kb | incomplete | 7 | 20935-25834 | PHAGE_Mycoba_Shipwreck_NC_031261(1) | 51.49% |
| pT115A | 1 | 11.6Kb | incomplete | 12 | 1-11645 | PHAGE_Escher_RCS47_NC_042128(3) | 51.70% |
| pT156A | 1 | 15.4Kb | incomplete | 12 | 2525-17988 | PHAGE_Escher_RCS47_NC_042128(4) | 50.37% |
| pT159A | 1 | 15.3Kb | incomplete | 26 | 310-15624 | PHAGE_Escher_RCS47_NC_042128(3) | 55.93% |
| pT199A | 1 | 11.6Kb | incomplete | 13 | 1-11638 | PHAGE_Escher_RCS47_NC_042128(3) | 51.70% |
| pT209A | 1 | 15.3Kb | incomplete | 26 | 266-15632 | PHAGE_Escher_RCS47_NC_042128(3) | 55.99% |
| pT224A | 1 | 17.9Kb | incomplete | 13 | 2525-20477 | PHAGE_Escher_RCS47_NC_042128(3) | 51.06% |
| pT247A | 1 | 20.5Kb | incomplete | 17 | 9428-29984 | PHAGE_Klebsi_phiKO2_NC_005857(2) | 55.10% |
|  | 2 | 9.1Kb | incomplete | 14 | 22591-31767 | PHAGE_Enterо_N15_NC_001901(2) | 56.61% |
| pT257A | 1 | 13.3Kb | incomplete | 8 | 2525-15842 | PHAGE_Escher_RCS47_NC_042128(4) | 50.74% |
| pT267A | 1 | 15.3Kb | incomplete | 26 | 266-15632 | PHAGE_Escher_RCS47_NC_042128(3) | 55.99% |
| pT270A | 1 | 7.1Kb | incomplete | 12 | 5753-12871 | PHAGE_Escher_RCS47_NC_042128(3) | 53.95% |
|  | 2 | 4.9Kb | incomplete | 7 | 20943-25842 | PHAGE_Mycoba_Shipwreck_NC_031261(1) | 51.49% |
| pT295A | 1 | 27.6Kb | incomplete | 21 | 3066-30726 | PHAGE_Escher_RCS47_NC_042128(3) | 48.94% |
| pT390A | 1 | 7.5Kb | incomplete | 13 | 52679-60238 | PHAGE_Cronob_ENT39118_NC_019934(2) | 41.87% |
|  | 2 | 33.7Kb | incomplete | 12 | 51310-85093 | PHAGE_Escher_RCS47_NC_042128(3) | 49.52% |
|  | 3 | 4.9Kb | incomplete | 7 | 79519-84418 | PHAGE_Mycoba_Shipwreck_NC_031261(1) | 51.49% |
| pT428A1 | 1 | 20.5Kb | incomplete | 17 | 9428-29984 | PHAGE_Klebsi_phiKO2_NC_005857(2) | 55.10% |
|  | 2 | 9.1Kb | incomplete | 14 | 22591-31767 | PHAGE_Enterо_N15_NC_001901(2) | 56.61% |
| pT415A | 1 | 16.6Kb | questionable | 20 | 1146-17798 | PHAGE_Salmon_SJ46_NC_031129(3) | 49.94% |
| pT455A | 1 | 9.3Kb | incomplete | 10 | 20175-29551 | PHAGE_Salmon_SJ46_NC_031129(4) | 53.36% |
| pT525A | 1 | 41Kb | questionable | 51 | 73364-114387 | PHAGE_Acinet_vB_AbaM_ME3_NC_041884(3) | 50.45% |
|  | 2 | 29.9Kb | questionable | 25 | 142809-172780 | PHAGE_Escher_RCS47_NC_042128(3) | 52.21% |
| pT526A | 1 | 24.6Kb | incomplete | 9 | 7334-31962 | PHAGE_Salmon_SJ46_NC_031129(4) | 53.01% |
| pT545A | 1 | 7.6Kb | incomplete | 12 | 391-7990 | PHAGE_Cronob_ENT39118_NC_019934(2) | 41.96% |
|  | 2 | 4.9Kb | incomplete | 7 | 81134-86033 | PHAGE_Mycoba_Shipwreck_NC_031261(1) | 51.49% |
|  | 3 | 7Kb | incomplete | 12 | 94105-101171 | PHAGE_Escher_RCS47_NC_042128(3) | 53.81% |
| pT570A | 1 | 24.6Kb | incomplete | 9 | 7371-31999 | PHAGE_Salmon_SJ46_NC_031129(4) | 53.01% |
| pT593A | 1 | 23.2Kb | incomplete | 14 | 1517-24805 | PHAGE_Escher_RCS47_NC_042128(3) | 53.30% |
|  | 2 | 6.4Kb | incomplete | 13 | 21185-27638 | PHAGE_Enterо_N15_NC_001901(2) | 55.38% |

The completeness was determined by PHASTER based on each hit score: Intact (score > 90), Questionable (score 70-90), Incomplete (score < 70).

There were seven most common phage detected on these input plasmids. PHAGE_Escher_RCS47_NC_042128 was detected in most of them (16/25) while others were less common on our plasmids. PHAGE_Mycoba_Shipwreck_NC_031261 was found on five plasmids. PHAGE_Entero_N15_NC_001901 and PHAGE_Salmon_SJ46_NC_031129 were found on four plasmids. PHAGE_Klebsi_phiKO2_NC_005857 and PHAGE_Cronob_ENT39118_NC_019934 were detected on three plasmids. PHAGE_Acinet_vB_AbaM_ME3_NC_041884 was only detected on the plasmid pT525A.

All input plasmids but two got prophage hits classified as incomplete with a score ≤ 70. The PHASTER tool has its own criteria for scoring prophage regions and classifying them based on their scores: intact (score > 90), questionable (score 70-90), incomplete (score < 70). Two plasmids got hits with scores in a range of 70-90 (questionable), higher scores compared to other plasmids (Fig. 4). These two plasmids included the largest plasmid pT525A with two hits and the IncY plasmid pT415A with one hit. None of the hits had the scores within the range of the intact group (>90).

**Fig. 4:**
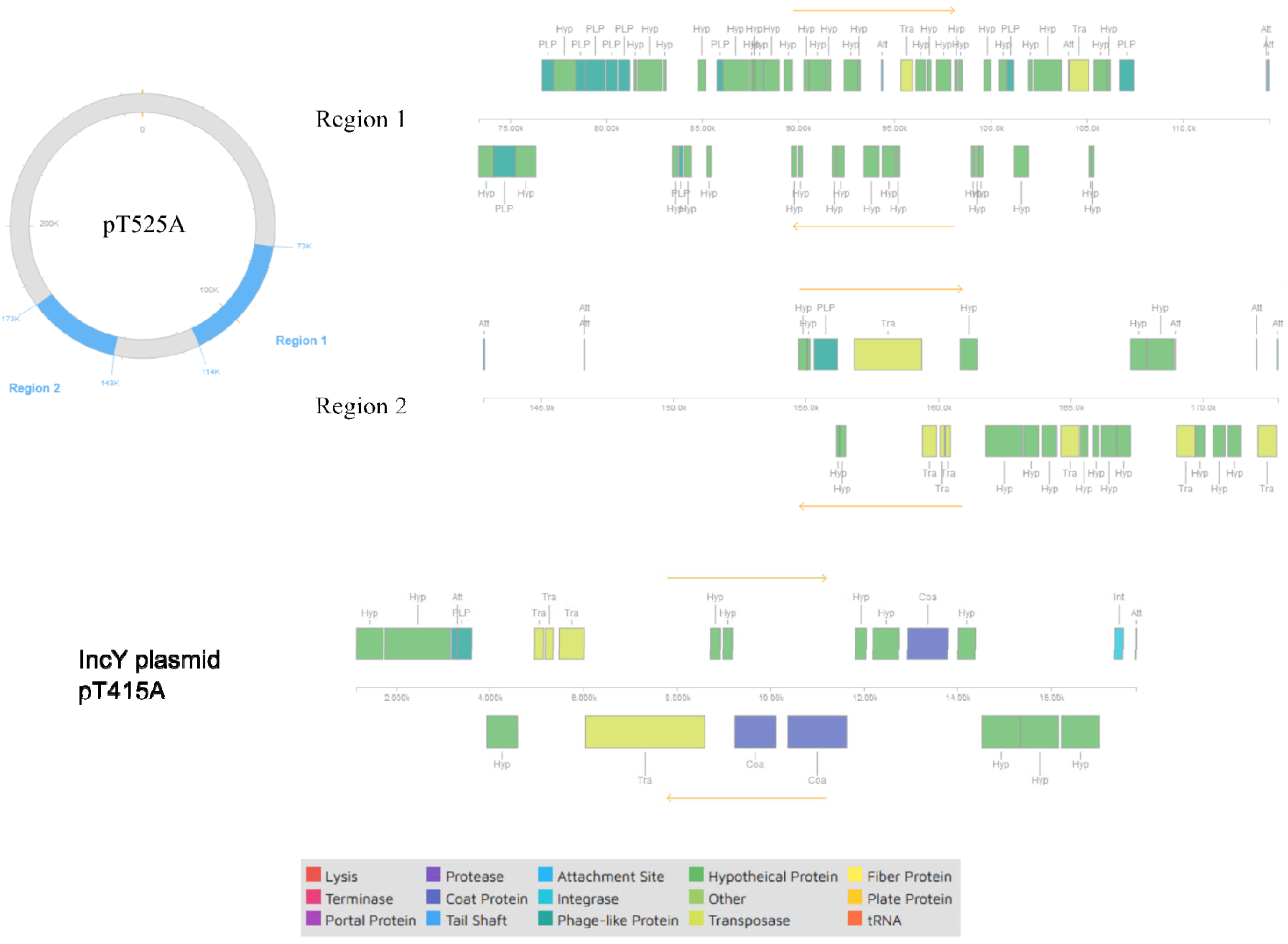
Map of putative prophage regions that were detected on plasmid pT525A (Region 1 & Region 2) and on IncY plasmid pT415A by PHASTER

## DISCUSSION

The most prevalent plasmid found among 83 sequenced plasmids was the 104 kb plasmid carrying the *bla*_CTXM-15_ gene. This result was consistent with our previous observation using restriction enzyme profiles combined with a subset of 25 sequenced plasmids (25). This plasmid was abundantly present in both raw and digested manures across all participating farms; however, the current bioinformatics tool was unable to identify its incompatibility group. This left us wondering if the plasmid adopted some novel incompatibility group which has not been recognized in the database. In a recently published study, the dominant ESBL plasmid isolated from 53 dairy farms located in southwest England was 220-kb IncHI2 plasmid carrying *bla*_CTX-M-32_ (26). This is interesting because only three out of 83 sequenced plasmids in our study belonged to IncHI2 group and had plasmid sizes larger than 200 kb; nevertheless, they carried *bla*_CTX-M-55_ gene along with other completely different resistance genes. In addition, SNP analysis on these three plasmids revealed a number of nucleotide modifications including deletion, insertion and

Sequencing another set of plasmids, the number of which was as twice as those in our previous study, allowed us to identify more plasmids that were less frequent (25). The second frequent plasmid, which was found in four out of six farms, especially quite dominant in farm 7, was the 43 kb plasmid of group IncN carrying the *bla*_CTX-M-1_ gene - the only resistance gene on this plasmid. In addition, plasmids from digested manure were more diverse than those from raw manure. Previously we showed that anaerobic digestion significantly reduced the conjugation frequency of ESBL-carrying plasmids in raw manure, but did not necessarily changed plasmid enzyme restriction profiles (25). The fact that a greater diversity of plasmids obtained from digested manure observed in this study can be explained as following. Firstly, it could be that the most frequent plasmid was overwhelming in raw manure, hence there was less chance for other plasmids to be selected for further analysis. Secondly, restriction enzyme profile combined with a small sequencing set did not adequately identify other less frequent plasmids. With a larger sequencing set, more distinct plasmids have been revealed in this study.

The *bla*_CMY-2_ gene is considered as an AmpC-type β-lactamase gene, which has been wide-spread around the world (27–29). This gene was mostly found on IncA/C plasmids in *E. coli* and *Salmonella* strains (27, 30, 31). Normally this gene confers AmpC resistance phenotype which hydrolyzes cephamycins as well as other third-generation cephalosporins and does not get inhibited by ESBL inhibitors (i.e. clavulanic acid, sulbactam, tazobactam) (32).  Previous studies showed that co-location of AmpC and ESBL genes resulted in a combined ESBL/AmpC phenotype (29, 33, 34). Detection of ESBL phenotype in a strain producing both AmpC and ESBL enzymes could be problematic because clavulanic acid, an ESBL inhibitor used in ESBL confirmatory tests, induces the high level expression of AmpC which, in turn, masks the synergy effects on ESBL (28, 33). However, in our study the *bla*_CMY-2_ gene found on IncI1 plasmid conferred ESBL phenotype in *E. coli* strain. Although ESBL-producing *E. coli* carrying only the *bla*_CMY-2_ gene was reported previously, there was not a clear explanation for this phenomenon (3). IncI1 plasmids carrying the *bla*_CMY-2_ gene were shown to widely spread among *E. coli* (35), and they shared a high degree of sequence similarity when isolated from Enterobacteriaceae with different epidemiological links (36). Unlike other plasmids carrying ESBL genes in this study, insertion sequences were not located proximal to the *bla*_CMY-2_ gene.

The *bla*_IMP-27_ gene, a novel metallo-β-lactamase, was firstly isolated in *Proteus mirabilis* in Ontario, Canada and presented in a conference in 2012 (37). Not until four years later, it was reported in published studies, including those found in unrelated *Proteus mirabilis* clinical isolates from two geographically distinct locations in the United States (38, 39). Another study showed that the gene was located on conjugative plasmids that was transferable from either *Proteus mirabilis* or *Providencia rettgeri* to *E. coli* (40). Not only was this gene found in clinical settings, but it was also recovered from the environment of a swine farrow-to-finish operation in the United States (41). The resistance phenotype to carbapenems and β-lactams conferred by this gene was quite distinct and might vary among host strains (38, 40). In our study, the *bla*_IMP-27_ gene, along with other resistance genes, was surprisingly found integrated into the host chromosome; while the conjugative plasmid isolated from this *E. coli* host was antibiotic resistance-free. We postulated that the integrated region originally got a ride on the plasmid, and it was then transferred to the host chromosome via homologous recombination as soon as the plasmid entered the host cell. To our best knowledge, this is the first time this gene was found in environmental samples in Canada.

TA systems, which were first detected on plasmids and later in bacterial chromosomes, play a vital role in plasmid stability as well as other positive roles in bacterial physiology, pathogenicity, and evolution (12, 42, 43). All TA systems detected in this study belonged to Type II TA systems which has been most extensively studied (12). TA systems were supposedly associated with stress-induced environment conditions which enabled stress-responsive proteases to degrade the antitoxin protein in Type II TA systems and free the toxin protein from the TA complex, resulting in cell growth inhibition or cell death (12, 14). The most frequent toxin-antitoxin system found in our study was the MazF-MazE system because this system was located on the most prevalent plasmid. Two TA systems detected in the study, MazF-MazE and RelE-RelB, belonged to super-families that were shown to be abundant and present on plasmids (42–44). Several TA systems, such as HipB-HipA or RelE-RelB, also caused the generation of persister cells, which went to a dormant state, in the presence of antibiotics, thus survived and became immune to antibiotics (9, 45–48).

Virulence factors detected on plasmids in this study were mostly related to colicin-producing genes (*cia*, *cib*) and the *traT* gene encoding outer membrane complement resistance protein. Colicins Ia and Ib are very similar structurally and able to absorb to common receptor sites on the bacterial outer membranes (49–51). Yet they do not share immunity specificity, hence cells are immune to either colicin Ia or Ib depending on which colicin gene they carry (49). Colicins inhibit protein and nucleic acid biosynthesis and uncouple electron transport from active transport, resulting in the loss of cellular potassium and magnesium which causes cell death (49). The virulence-associated non-conjugative plasmid carrying a *traT*-like gene was first identified in *Salmonella typhirium*; however, the *traT* gene was also found located within the transfer operon of conjugative F-like plasmids in *E. coli* (52, 53). The *traT* gene, one of two F cistrons, prevents the formation of cell contacts, and thus inhibits DNA transfer within the cell population (52). This gene is also needed for the resistance to serum bactericidal activity in *S. typhirium* and *E. coli* (53, 54). The *cia*, *cib* and *traT* genes were found on less frequent plasmids in this study, suggesting that these plasmids were limitedly accessible and only transferable among hosts of particular genetic backgrounds.

Polluted environments such as manured soil, animal farming, waste water and aquaculture can serve as a hot spot for co-selection of metal and antibiotic resistance (55–57). Evidences for metal-driven co-selection of multiple antibiotic resistance via co-resistance and cross-resistance mechanisms were well-documented (55, 56). Co-resistance mechanism occurred when metal/metalloid resistance genes were co-located with antibiotic resistance genes on the same plasmid (55). In this study, gene determinants for resistance to mercury (metal) and tellurium (metalloid), the *merAPTR* operon and the *terC* gene, were detected on two distinct conjugative multidrug resistant plasmids. The genetic linkage of mercury-and antibiotic-resistance genes on conjugative plasmids was demonstrated through mating between *Enterobacteriaceae* family bacteria and doubly genetically marked laboratory recipients (58). IncHI2 plasmids were known to be associated with tellurite resistance determinants previously (59, 60). Similarly, the *terC* gene was also detected on the largest IncHI2 plasmid in this study. Cross-resistance mechanism typically involved common efflux pump systems which pumped out structurally distinct agents/compounds such as metals and antibiotics (55, 56). Cheng et al. showed that chromosomally encoded TetA(L) efflux pump was able to remove both tetracycline and heavy metal cobalt (61). The *tet(A)* gene encoding major facilitator superfamily multidrug efflux pump was detected in several unrelated plasmids in this study including IncHI2, IncN, IncY, IncFIB(K) and IncI1 plasmids.

Prophage are normally found in bacterial chromosome, in particular within pathogenicity islands in pathogens (18, 62). A mounting number of studies showed that bacteriophages contributed to the widespread dissemination of antibiotic resistance genes via phage-mediated transduction (63, 64). In this study, PHASTER tool was used to investigate how likely prophage sequence could be detected on multi-drug resistant plasmids. Prophage regions were detected in a majority of input plasmids; however, the hit scores were pretty low, suggesting these prophages were unlikely to become active phages. The largest plasmid pT525A and the IncY plasmid pT425A had better scores for prophage hits. There was a possibility that phage was able to insert its sequences into plasmids, resulting in plasmids of larger size and more diversity as exemplified by plasmid pT525A. IncY plasmids were known to be phage-like plasmids because they shared a large portion of homologous segments with bacteriophage, in particular phage P1 (17, 19, 24, 65, 66).

In conclusion, this study genotypically characterized ESBL plasmids from dairy manure including plasmid sizes, antibiotic resistance genes, incompatibility groups, toxin-antitoxin systems, virulence factor and prophage regions. Sequencing a larger set of plasmids revealed more distinct less frequent plasmids, especially in digested manure. The *bla_IMP-27_* gene conferring resistance to both carbapenem and third-generation cephalosporis was found integrated into the host chromosome. The study provided some insights into the dynamics of ESBL genes and plasmids carrying these genes in dairy manure.

## MATERIALS AND METHODS

### 1. Conjugation method

The experiment was described in a previous study (25). Briefly, dairy manures were enriched with cefotaxime (4 mg/L) and then mated with *gfp-*labelled *E. coli* CV601 overnight. Transconjugants were selected on chromocult media containing rifampicin (50 mg/L), kanamycin (50 mg/L) and cefotaxime (4 mg/L).

### 2. Illumina/MinION sequencing protocol

#### 3. Annotation tools to detect antibiotic resistance genes, mobile genetic elements and toxin-antitoxin systems

Antibiotic resistance genes were detected using starAMR tool (Galaxy Version 0.7.1+galaxy1) which searched Illumina short-read assemblies against the resfinder resistance gene database. Mobile genetic elements were detected by RAST (https://rast.nmpdr.org), and then subsequently specified by blasting sequences against the NCBI non-redundant database. RAST was also used to detect toxin-antitoxin systems on complete closed plasmids.

#### 4. Detection of virulence factors

Complete closed plasmid sequences from hybrid assembly were used as input to VirulenceFinder 2.0, a web-tool (https://cge.cbs.dtu.dk/services/VirulenceFinder/), to detect virulence genes (67).

#### 5. Detection of prophage sequences

A web-tool PHASTER (http://phaster.ca/) was used to identify and annotate prophage sequences within complete closed plasmids (68, 69).

#### 6. Other tools used to construct plasmid maps and detect SNPs

## REFERENCE

1. Canton R, Gonzalez-Alba JM, Galan JC. 2012. CTX-M Enzymes: Origin and Diffusion. Front Microbiol 3:110.

2. Pitout JD, Laupland KB. 2008. Extended-spectrum β-lactamase-producing Enterobacteriaceae: an emerging public-health concern. Lancet Infect Dis 8:159–166.

3. Shin SW, Jung M, Won HG, Belaynehe KM, Yoon IJ, Yoo HS. 2017. Characteristics of Transmissible CTX-M-and CMY-Type beta-Lactamase-Producing Escherichia coli Isolates Collected from Pig and Chicken Farms in South Korea. J Microbiol Biotechnol 27:1716–1723.

4. Ben Said L, Jouini A, Klibi N, Dziri R, Alonso CA, Boudabous A, Ben Slama K, Torres C. 2015. Detection of extended-spectrum beta-lactamase (ESBL)-producing Enterobacteriaceae in vegetables, soil and water of the farm environment in Tunisia. Int J Food Microbiol 203:86–92.

5. Adelowo OO, Caucci S, Banjo OA, Nnanna OC, Awotipe EO, Peters FB, Fagade OE, Berendonk TU. 2018. Extended Spectrum Beta-Lactamase (ESBL)-producing bacteria isolated from hospital wastewaters, rivers and aquaculture sources in Nigeria. Environ Sci Pollut Res Int 25:2744–2755.

6. Jiang X, Cui X, Xu H, Liu W, Tao F, Shao T, Pan X, Zheng B. 2019. Whole Genome Sequencing of Extended-Spectrum Beta-Lactamase (ESBL)-Producing Escherichia coli Isolated From a Wastewater Treatment Plant in China. Front Microbiol 10:1797.

7. Gonggrijp MA, Santman-Berends I, Heuvelink AE, Buter GJ, van Schaik G, Hage JJ, Lam T. 2016. Prevalence and risk factors for extended-spectrum beta-lactamase-and AmpC-producing *Escherichia coli i*n dairy farms. J Dairy Sci 99:9001–9013.

8. Valentin L, Sharp H, Hille K, Seibt U, Fischer J, Pfeifer Y, Michael GB, Nickel S, Schmiedel J, Falgenhauer L, Friese A, Bauerfeind R, Roesler U, Imirzalioglu C, Chakraborty T, Helmuth R, Valenza G, Werner G, Schwarz S, Guerra B, Appel B, Kreienbrock L, Kasbohrer A. 2014. Subgrouping of ESBL-producing Escherichia coli from animal and human sources: an approach to quantify the distribution of ESBL types between different reservoirs. Int J Med Microbiol 304:805–16.

9. Huang CY, Gonzalez-Lopez C, Henry C, Mijakovic I, Ryan KR. 2020. hipBA toxin-antitoxin systems mediate persistence in Caulobacter crescentus. Sci Rep 10:2865.

10. Schureck MA, Maehigashi T, Miles SJ, Marquez J, Cho SE, Erdman R, Dunham CM. 2014. Structure of the Proteus vulgaris HigB-(HigA)2-HigB toxin-antitoxin complex. J Biol Chem 289:1060–70.

11. Thisted T, Sørensen NS, Wagner EG, Gerdes K. 1994. Mechanism of post-segregational killing: Sok antisense RNA interacts with Hok mRNA via its 5’-end singlestranded leader and competes with the 3’-end of Hok mRNA for binding to the mok translational initiation region. The EMBO journal 13:1960–1968.

12. Yamaguchi Y, Park JH, Inouye M. 2011. Toxin-antitoxin systems in bacteria and archaea. Annu Rev Genet 45:61–79.

13. Stieber D, Gabant P, Szpirer CY. 2008. The art of selective killing: plasmid toxin/antitoxin systems and their technological applications. BioTechniques 45:344–346.

14. Van Melderen L. 2010. Toxin-antitoxin systems: why so many, what for? Curr Opin Microbiol 13:781–5.

15. Johnson JR, Moseley SL, Roberts PL, Stamm WE. 1998. Aerobactin and Other Virulence Factor Genes among Strains of Escherichia coli Causing Urosepsis: Association with Patient Characteristics. Infection and immunity 56:405–412.

16. Wu HJ, Wang AH, Jennings MP. 2008. Discovery of virulence factors of pathogenic bacteria. Curr Opin Chem Biol 12:93–101.

17. Johnson TJ, Nolan LK. 2009. Pathogenomics of the virulence plasmids of Escherichia coli. Microbiol Mol Biol Rev 73:750–74.

18. Schmidt H, Hensel M. 2004. Pathogenicity islands in bacterial pathogenesis. Clin Microbiol Rev 17:14–56.

19. Frost LS, Leplae R, Summers AO, Toussaint A. 2005. Mobile genetic elements: the agents of open source evolution. Nat Rev Microbiol 3:722–32.

20. SONEA S. 1987. Bacterial Viruses, Prophages, and Plasmids, Reconsidered. Annals of the New York Academy of Sciences 503:251–260.

21. Wang F, Wang D, Hou W, Jin Q, Feng J, Zhou D. 2019. Evolutionary Diversity of Prophage DNA in Klebsiella pneumoniae Chromosomes. Front Microbiol 10:2840.

22. Casjens SR, Gilcrease EB, Huang WM, Bunny KL, Pedulla ML, Ford ME, Houtz JM, Hatfull GF, Hendrix RW. 2004. The pKO2 linear plasmid prophage of Klebsiella oxytoca. J Bacteriol 186:1818–32.

23. Czajkowski R. 2019. May the Phage be With You? Prophage-Like Elements in the Genomes of Soft Rot Pectobacteriaceae: Pectobacterium spp. and Dickeya spp. Front Microbiol 10:138.

24. Zhang C, Feng Y, Liu F, Jiang H, Qu Z, Lei M, Wang J, Zhang B, Hu Y, Ding J, Zhu B. 2017. A Phage-Like IncY Plasmid Carrying the *mcr-1* Gene in *Escherichia coli* from a Pig Farm in China. Antimicrob Agents Chemother 61.

25. Tran TT, Scott A, Tien YC, Murray R, Boerlin P, Pearl DL, Liu K, Robertson J, Nash JHE, Topp E. 2021. On-Farm Anaerobic Digestion of Dairy Manure Reduces the Abundance of Antibiotic Resistance-Associated Gene Targets and the Potential for Plasmid Transfer. Appl Environ Microbiol 87:e0298020.

26. Findlay J, Mounsey O, Lee WWY, Newbold N, Morley K, Schubert H, Gould VC, Cogan TA, Reyher KK, Avison MB. 2021. Molecular Epidemiology of Escherichia coli Producing CTX-M and pAmpC β-Lactamases from Dairy Farms Identifies a Dominant Plasmid Encoding CTX-M-32 but No Evidence for Transmission to Humans in the Same Geographical Region. Appl Environ Microbiol 87:e01842–20.

27. Guo YF, Zhang WH, Ren SQ, Yang L, Lu DH, Zeng ZL, Liu YH, Jiang HX. 2014. IncA/C plasmid-mediated spread of CMY-2 in multidrug-resistant Escherichia coli from food animals in China. PLoS One 9:e96738.

28. Botelho LA, Kraychete GB, Costa e Silva JL, Regis DV, Picao RC, Moreira BM, Bonelli RR. 2015. Widespread distribution of CTX-M and plasmid-mediated AmpC beta-lactamases in Escherichia coli from Brazilian chicken meat. Mem Inst Oswaldo Cruz 110:249–54.

29. Rizi KS, Mosavat A, Youssefi M, Jamehdar SA, Ghazvini K, Safdari H, Amini Y, Farsiani H. 2020. High prevalence of blaCMY AmpC beta-lactamase in ESBL co-producing Escherichia coli and Klebsiella spp. clinical isolates in the northeast of Iran. J Glob Antimicrob Resist 22:477–482.

30. Carattoli A, Villa L, Poirel L, Bonnin RA, Nordmann P. 2012. Evolution of IncA/C blaCMY-(2)-carrying plasmids by acquisition of the blaNDM-(1) carbapenemase gene. Antimicrob Agents Chemother 56:783–6.

31. Call DR, Singer RS, Meng D, Broschat SL, Orfe LH, Anderson JM, Herndon DR, Kappmeyer LS, Daniels JB, Besser TE. 2010. blaCMY-2-positive IncA/C plasmids from Escherichia coli and Salmonella enterica are a distinct component of a larger lineage of plasmids. Antimicrob Agents Chemother 54:590–6.

32. Hanson ND. 2003. AmpC beta-lactamases: what do we need to know for the future? J Antimicrob Chemother 52:2–4.

33. Shoorashetty RM, Nagarathnamma T, Prathibha J. 2011. Comparison of the boronic acid disk potentiation test and cefepime-clavulanic acid method for the detection of ESBL among AmpC-producing Enterobacteriaceae. Indian J Med Microbiol 29:297–301.

34. Smet A, Martel A, Persoons D, Dewulf J, Heyndrickx M, Catry B, Herman L, Haesebrouck F, Butaye P. 2008. Diversity of extended-spectrum beta-lactamases and class C beta-lactamases among cloacal Escherichia coli Isolates in Belgian broiler farms. Antimicrob Agents Chemother 52:1238–43.

35. Sidjabat HE, Seah KY, Coleman L, Sartor A, Derrington P, Heney C, Faoagali J, Nimmo GR, Paterson DL. 2014. Expansive spread of IncI1 plasmids carrying blaCMY-2 amongst Escherichia coli. Int J Antimicrob Agents 44:203–8.

36. Drijver E, Stohr J, Verweij JJ, Verhulst C, Velkers FC, Stegeman A, Bergh M, Kluytmans J, Group IS. 2020. Limited Genetic Diversity of blaCMY-2-Containing IncI1-pST12 Plasmids from Enterobacteriaceae of Human and Broiler Chicken Origin in The Netherlands. Microorganisms 8.

37. Tijet N, Lo S, Siebert H, MacNeill M, Rawte P, Farrell DJ, Low DE, Patel SN, Melano RG. 2012. Detection of IMP-27 metallo-β-lactamase in Proteus mirabilis, ON, Canada, abstr C2–090, Abstr 52nd Intersci Conf Antimicrob Agents Chemother.

38. Dixon N, Fowler RC, Yoshizumi A, Horiyama T, Ishii Y, Harrison L, Geyer CN, Moland ES, Thomson K, Hanson ND. 2016. IMP-27, a Unique Metallo-beta-Lactamase Identified in Geographically Distinct Isolates of Proteus mirabilis. Antimicrob Agents Chemother 60:6418–21.

39. Tijet N, Patel SN, Melano RG. 2016. Detection of carbapenemase activity in Enterobacteriaceae: comparison of the carbapenem inactivation method versus the Carba NP test. J Antimicrob Chemother 71:274–6.

40. Potter RF, Wallace MA, McMullen AR, Prusa J, Stallings CL, Burnham CAD, Dantas G. 2018. blaIMP-27 on transferable plasmids in Proteus mirabilis and Providencia rettgeri. Clin Microbiol Infect 24:1019 e5–1019 e8.

41. Mollenkopf DF, Stull JW, Mathys DA, Bowman AS, Feicht SM, Grooters SV, Daniels JB, Wittum TE. 2017. Carbapenemase-Producing Enterobacteriaceae Recovered from the Environment of a Swine Farrow-to-Finish Operation in the United States. Antimicrob Agents Chemother 61.

42. Leplae R, Geeraerts D, Hallez R, Guglielmini J, Dreze P, Van Melderen L. 2011. Diversity of bacterial type II toxin-antitoxin systems: a comprehensive search and functional analysis of novel families. Nucleic Acids Res 39:5513–25.

43. Gotfredsen M, Gerdes K. 1998. The Escherichia coli relBEgenes belong to a new toxin–antitoxin gene family. Molecular Microbiology 29:1065–1076.

44. Grønlund H, Gerdes K. 1999. Toxin-Antitoxin Systems Homologous with relBE of Escherichia coli Plasmid P307 are Ubiquitous in Prokaryotes. J Mol Biol 285:1401–1415.

45. Keren I, Shah D, Spoering A, Kaldalu N, Lewis K. 2004. Specialized persister cells and the mechanism of multidrug tolerance in Escherichia coli. J Bacteriol 186:8172–80.

46. Magnuson RD. 2007. Hypothetical functions of toxin-antitoxin systems. J Bacteriol 189:6089–92.

47. Korch SB, Henderson TA, Hill TM. 2003. Characterization of the hipA7 allele of Escherichia coli and evidence that high persistence is governed by (p)ppGpp synthesis. Mol Microbiol 50:1199–213.

48. Kussell E, Kishony R, Balaban NQ, Leibler S. 2005. Bacterial persistence: a model of survival in changing environments. Genetics 169:1807–14.

49. Mankovich JA, Hsu CH, Konisky J. 1986. DNA and amino acid sequence analysis of structural and immunity genes of colicins Ia and Ib. Journal of bacteriology 168:228–236.

50. Konisky J, Cowell BS. 1972. Interaction of Colicin Ia with Bacterial Cells. Journal of Biological Chemistry 247:6524–6529.

51. Konisky J, With the technical assistance of Billie SC. 1972. Characterization of Colicin Ia and Colicin Ib. Journal of Biological Chemistry 247:3750–3755.

52. Achtman M, Kennedy N, Skurray R. 1977. Cell-cell interactions in conjugating *Escherichia coli*: Role of traT protein in surface exclusion. Proc Nati Acad Sci USA 74:5104–5108.

53. Sukupolvi S, O’connor CD. 1990. TraT Lipoprotein, a Plasmid-Specified Mediator of Interactions between Gram-Negative Bacteria and Their Environment. MICROBIOLOGICAL REVIEWS 54:331–341.

54. Moll A, Manning PA, Timmis KN. 1980. Plasmid-Determined Resistance to Serum Bactericidal Activity: a Major Outer Membrane Protein, the traT Gene Product, Is Responsible for Plasmid-Specified Serum Resistance in *Escherichia coli*. INFECTION AND IMMUNITY 28:359–367.

55. Baker-Austin C, Wright MS, Stepanauskas R, McArthur JV. 2006. Co-selection of antibiotic and metal resistance. Trends Microbiol 14:176–82.

56. Imran M, Das KR, Naik MM. 2019. Co-selection of multi-antibiotic resistance in bacterial pathogens in metal and microplastic contaminated environments: An emerging health threat. Chemosphere 215:846–857.

57. Seiler C, Berendonk TU. 2012. Heavy metal driven co-selection of antibiotic resistance in soil and water bodies impacted by agriculture and aquaculture. Front Microbiol 3:399.

58. Summers AO, Wireman J, Vimy MJ, Lorscheider FL, Marshall B, Levy SB, Bennett S, Billard L. 1993. Mercury released from dental “silver” fillings provokes an increase in mercury-and antibiotic-resistant bacteria in oral and intestinal floras of primates. Antimicrobial Agents and Chemotherapy 37:825–834.

59. Hill SM, Jobling MG, Lloyd BH, Strike P, Ritchie DA. 1993. Functional expression of the tellurite resistance determinant from the IncHI-2 plasmid pMER610. Mol Gen Genet 241:203–12.

60. Whelan KF, Colleran E, Taylor DE. 1995. Phage inhibition, colicin resistance, and tellurite resistance are encoded by a single cluster of genes on the IncHI2 plasmid R478. Journal of Bacteriology 177:5016–5027.

61. Cheng J, Hicks DB, Krulwich TA. 1996. The purified Bacillus subtilis tetracycline efflux protein TetA(L) reconstitutes both tetracycline-cobalt/H+ and Na+(K+)/H+ exchange. Proceedings of the National Academy of Sciences of the United States of America 93:14446–14451.

62. Haaber J, Leisner JJ, Cohn MT, Catalan-Moreno A, Nielsen JB, Westh H, Penades JR, Ingmer H. 2016. Bacterial viruses enable their host to acquire antibiotic resistance genes from neighbouring cells. Nat Commun 7:13333.

63. Colavecchio A, Cadieux B, Lo A, Goodridge LD. 2017. Bacteriophages Contribute to the Spread of Antibiotic Resistance Genes among Foodborne Pathogens of the Enterobacteriaceae Family - A Review. Front Microbiol 8:1108.

64. Torres-Barcelo C. 2018. The disparate effects of bacteriophages on antibiotic-resistant bacteria. Emerg Microbes Infect 7:168.

65. Venturini C, Zingali T, Wyrsch ER, Bowring B, Iredell J, Partridge SR, Djordjevic SP. 2019. Diversity of P1 phage-like elements in multidrug resistant Escherichia coli. Sci Rep 9:18861.

66. Anonymous. <Sequence Relations among the IncY Plasmid ∼158, Pl , and P7 Prophages.pdf>.

67. Joensen KG, Scheutz F, Lund O, Hasman H, Kaas RS, Nielsen EM, Aarestrup FM. 2014. Real-time whole-genome sequencing for routine typing, surveillance, and outbreak detection of verotoxigenic *Escherichia coli*. J Clin Microbiol 52:1501–10.

68. Arndt D, Grant JR, Marcu A, Sajed T, Pon A, Liang Y, Wishart DS. 2016. PHASTER: a better, faster version of the PHAST phage search tool. Nucleic Acids Res 44:W16–21.

69. Zhou Y, Liang Y, Lynch KH, Dennis JJ, Wishart DS. 2011. PHAST: a fast phage search tool. Nucleic Acids Res 39:W347–52.

