## Supplementary file for "Genetic characterization of extended-β-lactamase (ESBL) plasmids captured from dairy manures"

Table:

| **ESBL gene** | **Representative plasmid** | **Other resistance genes co-locating on the same plasmid** | **Incompatibility group** | **Size** | **Farm** | **Raw**  **(n = 35)** | **Digested (n = 48)** |
| --- | --- | --- | --- | --- | --- | --- | --- |
| *bla*_CTX-M-1_ | pT115A | None | IncN | 42,592 | 1,2,5,7 | 10 | 8 |
| *bla*_CTX-M-1_ | pT39A | *sul2, tet(A)* | IncI1 | ~75,251 | 3 | NA | 1 |
| *bla*_CTX-M-14_ | pT428A | None | IncI1 | 91,905 | 2,3 | 3 | NA |
| *bla*_CTX-M-14_ | pT593A | None | IncB/O/K/Z | ~ 89,017 | 3 | NA | 1 |
| *bla*_CTX-M-14b_ | pT59A | *aac(6')-Ib-cr, aph(3'')-Ib, ARR-3,dfrA27, qnrS3, sul1, sul2, tet(A)* | IncFIB(K) |  | 5 | NA | 1 |
| *bla*_CTX-M-14b_ | pT58A | *aadA16, ARR-3, bla*_TEM-1B,_ *qnrS1, sul1, sul2, tet(A)* | IncFIB(K) |  | 5 | NA | 1 |
| *bla*_CTX-M-15_ | pT145A | None | IncI1 | ~ 85,051 | 1 | 2 | NA |
| *bla*_CTX-M-15_ | pT415A | *aph(3'')-Ib, aph(6)-Id, blaTEM-1B, dfrA14, qnrS1, sul2, tet(A)* | IncY | ~ 85,052 | 5 | NA | 1 |
| *bla*_CTX-M-15_ | pT545A | *aac(6')-Ib-cr, aph(3'')-Ib, aph(6)-Id, bla_OXA-1_, catB4, dfrA1, floR, lnu(G), sul2* | NA | 104,875 | 1,2,3,4,5,7 | 17 | 22 |
| *bla*_CTX-M-27_ | pT455A | None | IncN | 42,273 | 2 | NA | 1 |
| *bla*_CTX-M-27_ | pT257A | None | IncFIIA,IncFII | 66,581 | 2 | NA | 2 |
| *bla*_CTX-M-27_ | pT570A | *aph(3'')-Ib, aph(6)-Id, blaTEM-1B, dfrA14, sul2, tet(A)* | IncN | 53,066 | 2 | NA | 3 |
| *bla*_CTX-M-27_ | pT295A | *aph(3'')-Ib, aph(6)-Id, blaTEM-1B, dfrA14, sul2, tet(A)* | IncN, IncX1 | 77,311 | 2 | NA | 1 |
| *bla*_CTX-M-55_ | pT525A | *aac(3)-IId, aadA22, aph(3')-Ia, ARR-2, blaTEM-1B, dfrA14, floR, lnu(F), mph(A), qnrS1, sul3, tet(A)* | IncHI2, IncHI2A | 266,763 | 3 | 2 | NA |
| *bla*_CTX-M-55_ | pT588A | *aac(3)-IId, aadA22, aph(3')-Ia, ARR-2, bla*_TEM-1B_*, dfrA14, floR, lnu(F), mph(A), qnrS1, sul3, tet(A)* | IncHI2, IncHI2A, IncN | ~ 227,236 | 7 | NA | 1 |
| *bla*_CTX-M-55_ | pT476A | *blaTEM-1B* | IncX1 | ~ 40,307 | 7 | NA | 1 |
| *bla*_CTX-M-55_ | pT156A | *blaTEM-206* | IncFII(pHN7A8) | 66,894 | 1 | NA | 1 |
| *bla*_CTX-M-55_ | pT224A | *blaTEM-206, fosA3* | IncFII(pHN7A8) | ~ 70,625 | 7 | NA | 2 |
| *bla*_PER-1_ | pT413A | *aadA2, aph(3')-Ia, mph(E), msr(E), sul1, sul1, tet(C), tet(E), tet(X)* | IncC | 187,012 | 5 | 1 | NA |
| *bla*_IMP-27_ | pT598A | *aph(6)-Id, strA, sul2, tet(A)* | NA | 41,847 | 4 | NA | 1 |

Table 1: SNPs found on *bla*_CTX-M-1_-bearing plasmids when mapping Illumina sequence reads from other transconjugants’genomes against a reference genome (T115A) using SNIPPY

| **Farm 5/ Raw** | | | | **Farm 7/ Raw** | | | | **Farm 7/ Digestate With** | | | | **Farm 7/ Raw** | | | |
| --- | --- | --- | --- | --- | --- | --- | --- | --- | --- | --- | --- | --- | --- | --- | --- |
| **T110A** | | | | **T314A** | | | | **T324A** | | | | **T368A** | | | |
| POS | TYPE | REF | ALT | POS | TYPE | REF | ALT | POS | TYPE | REF | ALT | POS | TYPE | REF | ALT |
| N/D | | | | N/D | | | | N/D | | | | 30913 | snp | T | C |
| **Farm 7/ Digestate With** | | | | **Farm 7/ Raw** | | | | **Farm 7/ Digestate With** | | | | **Farm 7/ Digestate With** | | | |
| **T468A** | | | | **T548A** | | | | **T558A** | | | | **T324A** | | | |
| POS | TYPE | REF | ALT | POS | TYPE | REF | ALT | POS | TYPE | REF | ALT | POS | TYPE | REF | ALT |
| N/D | | | | N/D | | | | N/D | | | | N/D | | | |
|  |  |  |  |  |  |  |  | 39454 | snp | A | T |  |  |  |  |
| **Farm 1/ Digestate With** | | | | **Farm 7/ De-watered** | | | | **Farm 7/ Digestate With** | | | | **Farm 7/ Raw** | | | |
| **T486A** | | | | **T344A** | | | | **T378A** | | | | **T438A** | | | |
| POS | TYPE | REF | ALT | POS | TYPE | REF | ALT | POS | TYPE | REF | ALT | POS | TYPE | REF | ALT |
| N/D | | | | N/D | | | | N/D | | | | N/D | | | |
| **Farm 7/ Digestate W/O** | | | | **Farm 2/ Raw** | | | | **Farm 7/ Raw** | | | | **Farm 7/ Raw** | | | |
| **T451A** | | |  | **T599A** | | | | **T459A** | | | | **T578A** | | | |
| POS | TYPE | REF | ALT | POS | TYPE | REF | ALT | POS | TYPE | REF | ALT | POS | TYPE | REF | ALT |
| N/D | | | | 3222 | ins | A | ATTTTGCCACCCTAT | N/D | | | | N/D | | | |
|  |  |  |  | 39454 | snp | A | T |  |  |  |  |  |  |  |  |

Table 2: SNPs found on *bla*_CTX-M-15_-bearing plasmids when mapping Illumina sequence reads from other transconjugants’genomes against a reference genome (T159A) using SNIPPY

| **Farm 3 / Digestate With** | | | | **Farm 3 / Raw** | | | | **Farm 3/ Digestate W/O** | | | | | | **Farm 3 / Digestate W/O** | | | |
| --- | --- | --- | --- | --- | --- | --- | --- | --- | --- | --- | --- | --- | --- | --- | --- | --- | --- |
| **T1A** | | | | **T38A** | | | | **T48A** | | | | | | **T49A** | | | |
| POS | TYPE | REF | ALT | POS | TYPE | REF | ALT | POS | TYPE | | REF | | ALT | POS | TYPE | REF | ALT |
| N/D | | | | N/D | | | | N/D | | | | | | N/D | | | |
| **Farm 1/ Raw** | | | | **Farm 1/ Digestate with** | | | | **Farm 7/ Digestate With** | | | | | | **Farm 3/ Digestate With** | | | |
| **T63A** | | | | **T72A** | | | | **T125A** | | | | | | **T173A** | | | |
| POS | TYPE | REF | ALT | POS | TYPE | REF | ALT | POS | TYPE | | REF | | ALT | POS | TYPE | REF | ALT |
| 35405 | snp | A | G | 35405 | snp | A | G | 9489 | ins | | A | | ACGCCGGCGCGCATCGTCGGATGTGGCGATGCGAGCTCGAGGTCGGTGGCCAC | N/D | | | |
| **Farm 3/ Digestate W/O** | | | | **Farm 4/ Raw** | | | | **Farm 4/ Digestate With** | | | | | | **Farm 5/ Raw** | | | |
| **T180A** | | | | **T189A** | | | | **T192A** | | | | | | **T195A** | | | |
| POS | TYPE | REF | ALT | POS | TYPE | REF | ALT | POS | TYPE | | REF | | ALT | POS | TYPE | REF | ALT |
| N/D | | | | 35405 | snp | A | G | 35405 | snp | | A | | G | 36942 | snp | G | T |
| **Farm 7/ Digestate W/O** | | | | **Farm 7/ De-watered** | | | | **Farm 7/ De-watered** | | | | | | **Farm 7/ Heat Trt Compo** | | | |
| **T218A** | | | | **T228A** | | | | **T231A** | | | | | | **T238A** | | | |
| POS | TYPE | REF | ALT | POS | TYPE | REF | ALT | POS | TYPE | | REF | | ALT | POS | TYPE | REF | ALT |
| N/D | | | | N/D | | | | N/D | | | | | | N/D | | | |
| **Farm 5/ Raw** | | | | **Farm 7/ Digestate With** | | | | **Farm 4/ Raw** | | | | | | **Farm 4/ Raw** | | | |
| **T390A** | | | | **T442A** | | | | **T506A** | | | | | | **T508A** | | | |
| POS | TYPE | REF | ALT | POS | TYPE | REF | ALT | POS | TYPE | | REF | | ALT | POS | TYPE | REF | ALT |
| N/D | | | | N/D | | | | 35405 | snp | A | | G | | 35405 | snp | A | G |
| **Farm 4/ Digestate With** | | | | **Farm 4/ Digestate With** | | | | **Farm 3/ Digestate With** | | | | | | **Farm 2/ Raw** | | | |
| **T516A** | | | | **T523A** | | | | **T535A** | | | | | | **T545A** | | | |
| POS | TYPE | REF | ALT | POS | TYPE | REF | ALT | POS | TYPE | | REF | | ALT | POS | TYPE | REF | ALT |
| 35405 | snp | A | G | 35405 | snp | A | G | N/D | | | | | | N/D | | | |

Table 3: SNPs found on *bla*_CTX-M-27_-bearing plasmids when mapping Illumina sequence reads from other transconjugants’genomes against a reference genome (T570A) using SNIPPY

| **Farm 2/ Raw** | | | | **Farm 2/ Digestate With** | | | | **Farm 2/ Digestate With** | | | |
| --- | --- | --- | --- | --- | --- | --- | --- | --- | --- | --- | --- |
| **T31A** | | | | **T358A** | | | | **T192A** | | | |
| POS | TYPE | REF | ALT | POS | TYPE | REF | ALT | POS | TYPE | REF | ALT |
| N/D | | | | 46156 | ins | C | CG | 41414 | snp | T | C |

Table 4: SNPs found on *bla*_CTX-M-55_-bearing plasmids (> 200 kb) when mapping Illumina sequence reads from other transconjugants’genomes against a reference genome (T525A) using SNIPPY

| **Farm 7/ Digested** | | | | **Farm 3/ Raw** | | | |
| --- | --- | --- | --- | --- | --- | --- | --- |
| **T588A** | | | | **T594A** | | | |
| POS | TYPE | REF | ALT | POS | TYPE | REF | ALT |
| 17773 | del | CA | C | 10402 | snp | C | T |
| 18185 | del | CA | C | 17078 | complex | G | CA |
| 19480 | ins | C | CT | 17773 | del | CA | C |
| 20365 | ins | C | CT | 17850 | ins | G | GT |
| 49573 | ins | C | CTT | 18185 | del | CA | C |
| 49588 | ins | A | AC | 19480 | ins | C | CT |
| 49778 | del | AAG | A | 20365 | ins | C | CT |
| 49878 | ins | G | GA | 49573 | ins | C | CTT |
| 76461 | ins | G | GT | 49588 | ins | A | AC |
| 93495 | ins | C | CT | 49778 | del | AAG | A |
| 94375 | snp | G | A | 49950 | ins | G | GA |
| 94389 | ins | C | CAG | 49956 | ins | C | CT |
| 96661 | ins | G | GA | 94375 | snp | G | A |
| 97173 | ins | C | CCA | 94389 | ins | C | CAG |
| 97893 | ins | G | GA | 94417 | ins | C | CT |
| 98111 | ins | C | CT | 96661 | ins | G | GA |
| 98229 | ins | T | TA | 96879 | ins | G | GA |
| 154015 | ins | T | TC | 97893 | ins | G | GA |
| 154435 | ins | G | GA | 98111 | ins | C | CT |
| 155232 | ins | T | TA | 98229 | ins | T | TA |
| 177771 | ins | T | TG | 98437 | ins | T | TC |
| 221140 | snp | G | T | 154015 | ins | T | TC |
| 226044 | del | TA | T | 154435 | ins | G | GA |
|  |  |  |  | 155232 | ins | T | TA |
|  |  |  |  | 177771 | ins | T | TG |
|  |  |  |  | 189505 | ins | A | AT |
|  |  |  |  | 221140 | snp | G | T |
|  |  |  |  | 226044 | del | TA | T |
